# Systematic evaluation of *Cyanidioschyzon merolae* across photobioreactor systems: Linking reactor design to biomass production and biochemical composition

**DOI:** 10.64898/2026.06.17.732901

**Authors:** Philipp Ernst, Jan Vanselow, Max Denter, Wen Li, Lennart Witting, Jochem Gätgens, Markus Pauly, Dietrich Kohlheyer, Vlada B. Urlacher, Michael Feldbrügge, Julia Frunzke

## Abstract

Extremophilic red microalgae are promising platforms for sustainable biotechnology, combining robust growth under selective thermoacidophilic conditions with production of thermostable phycobiliproteins and carbon-rich biomass. However, reactor-dependent effects on growth, product formation and biomass composition remain insufficiently resolved. Here, we systematically evaluated the extremophilic red microalga *Cyanidioschyzon merolae* across cultivation scales and reactor formats and benchmarked its performance against the well-established *Galdieria javensis* and *Limnospira platensis*. In small-scale multi-cultivator photobioreactors and microfluidic growth chambers, *C. merolae* showed superior growth, reaching a maximum growth rate of 0.034 ± 0.001 h^-1^ and 8.3 ± 0.3 g l^-1^ cell dry weight. Microfluidic cultivation enabled growth analysis at single-cell resolution and matched growth rates obtained in photobioreactors. To identify scalable production strategies, *C. merolae* was further cultivated in a flat-panel photobioreactor and a custom-designed internally illuminated photobioreactor. The custom-designed photobioreactor delivered the highest biomass concentration and productivity, yielding 11.5 ± 0.6 g l^-1^ cell dry weight and 1.07 ± 0.06 g l^-1^ d^-1^, and comparable yields with regard to R-phycocyanin and R-allophycocyanin. Biomass analysis revealed substantial carbon and nitrogen contents, starch accumulation up to > 20 % of cell dry weight, and fatty acids dominated by palmitic, linoleic and oleic acids. Despite its reduced cell wall fraction, *C. merolae* contained structurally diverse, cultivation-dependent polysaccharides. These results establish *C. merolae* as a versatile chassis for thermostable pigment production and renewable feedstock generation, highlighting photobioreactor design as a key determinant of productivity and biomass quality.

## Introduction

With rising carbon dioxide (CO_2_) concentrations in the atmosphere, the need for efficient carbon capture methods is becoming increasingly important in counteracting climate change (Jones et al., 2024). Among others, biological carbon capture approaches using algae have gained much attention. Algae fix atmospheric CO_2_ through photosynthesis, require less space and exhibit higher growth rates than terrestrial plants (Ashour et al., 2024; Krause-Jensen & Duarte, 2016). In addition to their role in CO_2_ fixation, algae are gaining more and more interest for the production of valuable products. A prominent example is the cyanobacterium *Limnospira platensis* PCC7345, formerly known as *Spirulina platensis*, which is the current industrial producer of the blue, water-soluble protein C-phycocyanin (C-PC), a component of the light-harvesting complex (Ashour et al., 2024; Hamed, 2016; Mao et al., 2024). C-PC is widely used in the food and cosmetic industry, and is also studied in the medical and pharmaceutical context for its anti-oxidant, anti-inflammatory, anti-diabetic and anti-tumor properties (Mao et al., 2024). However, *L. platensis* is challenging to cultivate due to its unique life cycle and filamentous cell morphology. The relatively low thermostability of its C-PC further limits broader application. Several strategies have been developed to overcome the low thermostability, such as the addition of stabilising agents or the microencapsulation of C-PC (Mao et al., 2024). As these approaches are typically associated with higher costs, usage of more thermostable phycobiliproteins from other species represents a promising alternative, including R-phycocyanin (R-PC) from *rhodophyta* such as *Cyanidioschyzon merolae* 10D (Graziani et al., 2013; Rahman et al., 2017; Retta et al., 2024; Wan et al., 2016; Yoshida et al., 2021) and *Galdieria javensis* 074G (former *G. sulphuraria* 074G (Ott & Seckbach, 1994)).

*C. merolae* and *G. javensis* are unicellular thermoacidophilic red microalgae that synthesize and store large quantities of floridean starch in cytoplasmic granules during photosynthesis. Floridean starch is a glycogen-like storage polysaccharide consisting primarily of α-(1,4)-linked glucan chains interconnected by α-(1,6)-linked branch points (Pancha & Imamura, 2026). Its potential use as a carbon source for heterotrophic microorganisms highlights the utilization of these red microalgae not just as promising source of several compounds, but also as a sustainable alternative feedstock for biotechnological applications. However, a known bottleneck in the utilization of intracellular storage compounds from red microalgae is the resilience of their cells to disruption. Notably, *C. merolae* and *G. javensis* differ substantially in their cellular architecture, which leads to distinct downstream processing characteristics. Whereas *G. javensis* possesses a thick cell wall that complicates cell disruption and product recovery (Ashour et al., 2024; Retta et al., 2024; Wan et al., 2016), *C. merolae* lacks a rigid cell wall, which facilitates genetic manipulation and improves access to intracellular products (Rahman et al., 2017). This feature offers pronounced advantages for downstream processing.

The development of sustainable and easily processable carbon sources is of increasing importance, as conventional substrates such as glucose are typically derived from agricultural feedstocks and are associated with land use and additional CO_2_ emissions. In contrast, microalgae are able to convert CO_2_ into carbohydrates more efficiently than plants through photosynthesis (Singh et al., 2019), while simultaneously avoiding competition with food crop production. Importantly, red microalgae can even be cultivated in wastewater, enabling simultaneous nutrient removal (nitrogen, phosphorus and carbon assimilation) and biomass production. This supports a circular bioeconomy by coupling wastewater remediation with generation of valuable biomass as feedstock for downstream heterotrophic processes (Singh et al., 2023).

In this study, we systematically assessed the biotechnological potential of *C. merolae* by comparing its growth characteristics with those of *G. javensis* and *L. platensis* in small-scale multi-cultivator photobioreactors (MC) and microfluidic growth chambers. Furthermore, we benchmarked the standard MC photobioreactor against larger, scalable configurations, including a custom-designed internal-illuminated photobioreactor (IIPBR) and a flat-panel photobioreactor (FPR), to identify suitable cultivation strategies for maximizing *C. merolae* biomass productivity and yield. Finally, the resulting biomass was comprehensively analyzed with respect to pigment and fatty acid content, as well as elemental and monosaccharide composition, highlighting the advantages of *C. merolae* as a versatile platform organism for biotechnology and its potential as a renewable feedstock for heterotrophs.

## Results and Discussion

### *C. merolae* shows better growth performance compared to *G. javensis* and *L. platensis* in both multi-cultivator photobioreactors and microfluidic growth chambers

*C. merolae* was compared to *G. javensis* and *L. platensis* with respect to their growth performance in small-scale multi-cultivator reactors (Figure 1) and microfluidic growth chambers (Figure 2). If not otherwise stated, all cultivations were performed under standard conditions with 5 % CO_2_ (v v^-1^) and 400 µmol photos m^-2^ s^-1^ (µE).

**Figure 1:**
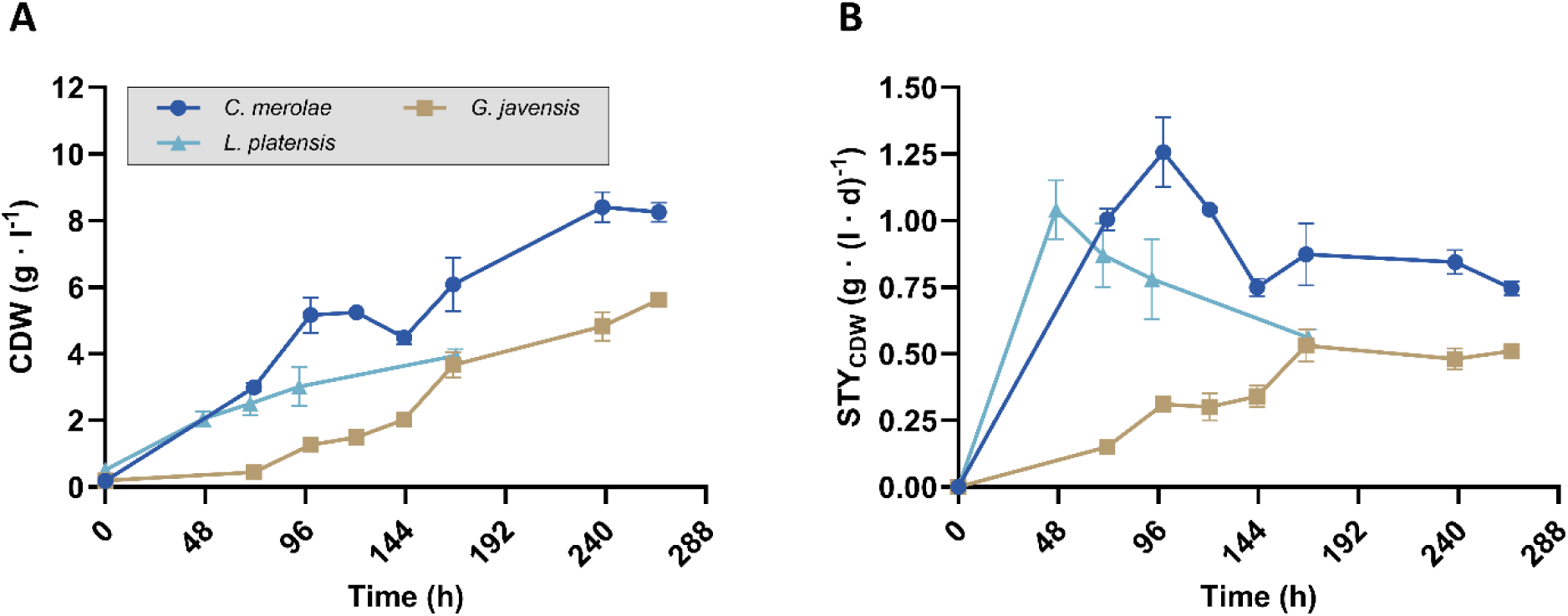
Growth comparison of *C. merolae*, *G. javensis* and *L. platensis* in multi-cultivators. (A) Cell dry weight (CDW) calculated form optical density (OD) values at 750 nm (Figure S1A and B) and from OD values at 680 nm for *L. platensis* (Figure S1C). (B) Space-time-yield CDW (STY_CDW_) during the cultivations, calculated from the CDW values and cultivation times (Table 1). Cultivations were performed at 42 °C for *C. merolae* and *G. javensis* and at 30 °C for *L. platensis*. The cultures were continuously aerated with 10.6 standard liter per hour (sl h^-1^) of 5 % CO_2_ and illuminated with 400 µE. *C. merolae* was grown in MA2 Medium, *G. javensis* in 2xGS Modified Allen Medium and *L. platensis* in Spirulina Medium. *L. platensis* had already reached the stationary phase after 168 h (n=3).

**Figure 2:**
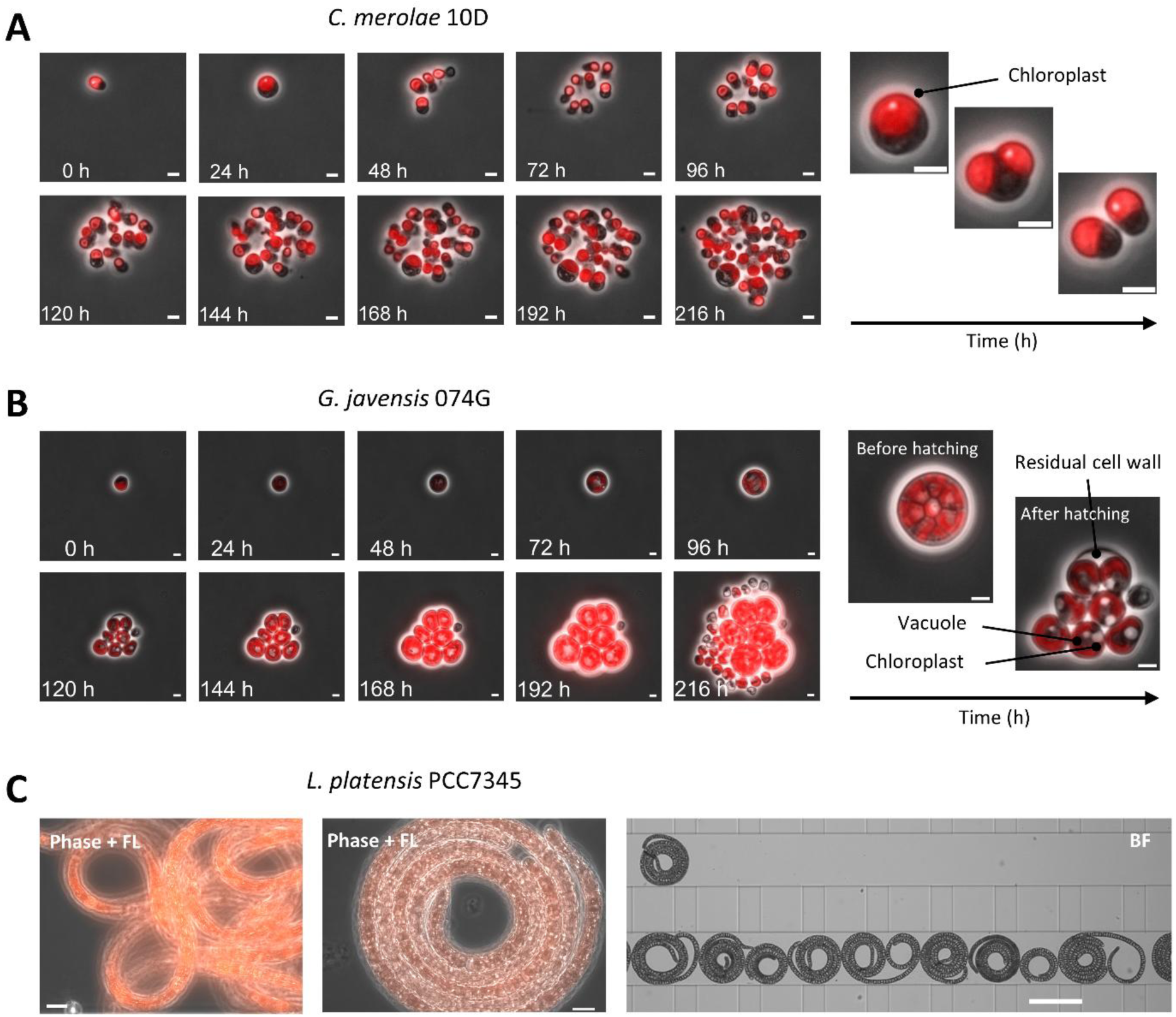
Growth of *C. merolae*, *G. javensis* and *L. platensis* in microfluidic growth chambers. Cultivations were performed at 42 °C for *C. merolae* and *G. javensis* and at 30 °C for *L. platensis*. The cultures were continuously illuminated with 30 µE under ambient CO_2_ conditions. *C. merolae* was grown in MA2 Medium, *G. javensis* in 2xGS Modified Allen Medium and *L. platensis* in Spirulina Medium. Scale bars represent 5 µm in panels A and B. Scale bars in the corresponding shorter time-lapse images represent 2 µm (A) and 3 µm (B). In panel C, scale bars represent 10 µm in the first and second images and 100 µm in the third image. A higher temporal resolution of microalgae growth dynamics is provided in Video S1 and S2.

**Table 1:** Growth comparison of *C. merolae*, *G. javensis* and *L. platensis* cultivations in the multi-cultivator and microfluidic growth chamber. Cultivations were performed at 42 °C for *C. merolae* and *G. javensis* and at 30 °C for *L. platensis*. The cultures were continuously aerated with 10.6 sl h^-1^ of 5 % CO_2_ and illuminated with 400 µE. *C. merolae* was grown in MA2 Medium, *G. javensis* in 2xGS Modified Allen Medium and *L. platensis* in Spirulina Medium. The mean values with standard deviation of three independent biological replicates are shown (n=3). For cultivations in microfluidic chips, identical temperatures were applied, while CO_2_ concentrations and light intensities were adjusted to single-layer cell growth.

| Parameter | Multi-cultivator |  |  | Microfluidics |  |
| --- | --- | --- | --- | --- | --- |
|  | *C. merolae* | *G. javensis* | *L. platensis* * | *C. merolae* | *G. javensis* |
| t (h) | 266 | 266 | 168 | 96 | 216 |
| $\mu_{\text{max}} (\text{h}^{-1})$ | $0.034 \pm 0.001$ | $0.022 \pm 0.001$ | $0.029 \pm 0.002$ | $0.033 \pm 0.001$ | $0.023 \pm 0.000$ |
| $t_d (\text{h})$ | $20.4 \pm 0.9$ | $31.4 \pm 0.2$ | $23.9 \pm 1.9$ | $20.7 \pm 0.5$ | $30 \pm 0.5$ |
| Final OD (-) | $28.8 \pm 1$ | $18.8 \pm 0.4$ | $3.9 \pm 0.2$ | | |
| Final CDW ($\text{g} \cdot \text{L}^{-1}$) | $8.3 \pm 0.3$ | $5.6 \pm 0.1$ | $3.9 \pm 0.2$ | | |
| $\text{STY}_{\text{CDW}} (\text{g} \cdot (\text{L} \cdot \text{d})^{-1})$ | $0.75 \pm 0.03$ | $0.51 \pm 0.02$ | $0.56 \pm 0.02$ | | |

\*Not in microfluidics.

As shown in Figure 1A and Table 1, *C. merolae* exhibited the most favorable performance among the three strains. With a growth rate of 0.034 ± 0.001 h^-1^ and a doubling time of 20.4 ± 0.9 h, it outcompeted *G. javensis* by a factor of almost 2. This was coupled to the highest final optical density (28.8 ± 1) and the highest final CDW (8.3 ± 0.3 g L^-1^) among the three investigated strains. As shown in Figure 1B, *C. merolae* also achieved the highest overall STY_CDW_ of 0.75 ± 0.03 g l^-1^ d^-1^ after 266 h, although *C. merolae* even achieved significantly higher STY_CDW_ of up to 1.25 g L^-1^ h^-1^ after around 100 h. Compared to previous reports, this is comparatively high for microalgal cultivations (McGinn et al., 2016; Seger et al., 2023). *G. javensis* reached an overall STY_CDW_ of 0.51 ± 0.02 g L^-1^ h^-1^ after 266 h, which is significantly lower than the value obtained for *C. merolae*, due to its slower growth rate of 0.022 ± 0.001 h^-1^. Additionally, *G. javensis* displayed a notably longer lag phase of approximately 70 h. The lowest final CDW was observed for *L. platensis* (3.9 ± 0.2 g l^-1^) as this strain entered stationary phase already after around 150 h of cultivation (Figure 1A).

*C. merolae* was further characterized with respect to varying CO_2_ concentrations, light intensities and light regimes, as these parameters strongly influence growth (Figure S2). Cultivation experiments in the MC revealed that growth rates were optimal under standard conditions of 5 % CO_2_ and 400 µE, whereas lower light intensities reduced growth and higher intensities caused saturation effects. Growth at 1 % CO_2_ was consistently lower, while 10 % CO_2_ resulted in extended lag phases and oscillatory growth behavior. In addition, light/dark cycles (14 h light / 10 h dark) negatively affected biomass accumulation, with decreases in OD_720_ observed during dark phases, whereas continuous illumination supported more stable and efficient growth, consistent with previous reports (Seger et al., 2023).

The three strains were additionally cultivated in microfluidic growth chambers (1.72 µm height) to visualize their division behavior and precisely determine their growth rates at single-cell resolution, enabling a more exact and time-resolved analysis than the bioreactor-based multi-cultivator experiments (Figure 2). *C. merolae* proliferates by binary fission (Krupnik et al., 2023), which was also observed in the microfluidic growth chambers (Figure 2A). Its simple cellular organization, characterized by the presence of only one organelle of each type (e.g., a single chloroplast and mitochondrion), was likewise confirmed. Notably, *C. merolae* exhibited morphological changes during growth, displaying a more irregular, fluid-like shape during cell division and returning to a round morphology upon completion. In contrast, *G. javensis* maintained a spherical morphology throughout cultivation and reproduced by successive cell divisions, forming 4 to 32 daughter cells within a mother cell (Figure 2B). Microscopic images in Figure 2B illustrate a single progenitor cell before and after hatching. *L. platensis* was also grown in a microfluidic chip (Figure 2C), but their size and morphology prevented reliable growth rate quantification from the recorded videos.

For *C. merolae* and *G. javensis*, growth rates determined from microfluidic experiments were in good agreement with those obtained from the multi-cultivator cultivations (Table 1). In addition, to the best of our knowledge, this study represents the first microfluidic growth characterization of *C. merolae*, *G. javensis* and *L. platensis*.

Overall, *C. merolae* exhibited the most promising growth characteristics among the tested strains. Its favorable growth performance is further supported by the absence of a rigid cell wall, which substantially facilitates downstream pigment extraction as no mechanical cell disruption is required.

### *C. merolae* yields the highest biomass concentration in internally illuminated photobioreactor

As efficient cultivation platforms are essential for the industrial exploitation of extremophilic microalgae, the MC system was benchmarked against different larger, scalable photobioreactor platforms. Reactor design strongly influences biomass and pigment productivity, as well as operational cost, making the selection of a suitable system critical for process development (Borowitzka, 2013; Carvalho et al., 2006; Posten, 2009). Various photobioreactor configurations are available, including vertical column systems such as the multi-cultivator, internal-illuminated photobioreactors, in which additional light is distributed directly into the culture via immersed glass rods, as well as flat-panel photobioreactors, which are characterized by a high surface-to-volume ratio and enhanced light availability.

*C. merolae* was cultivated in all photobioreactor setups under standard conditions at 42 °C and 5 % CO_2_ (Figure 3). Light intensities were adjusted according to the specific requirements of each cultivation system.

**Figure 3:**
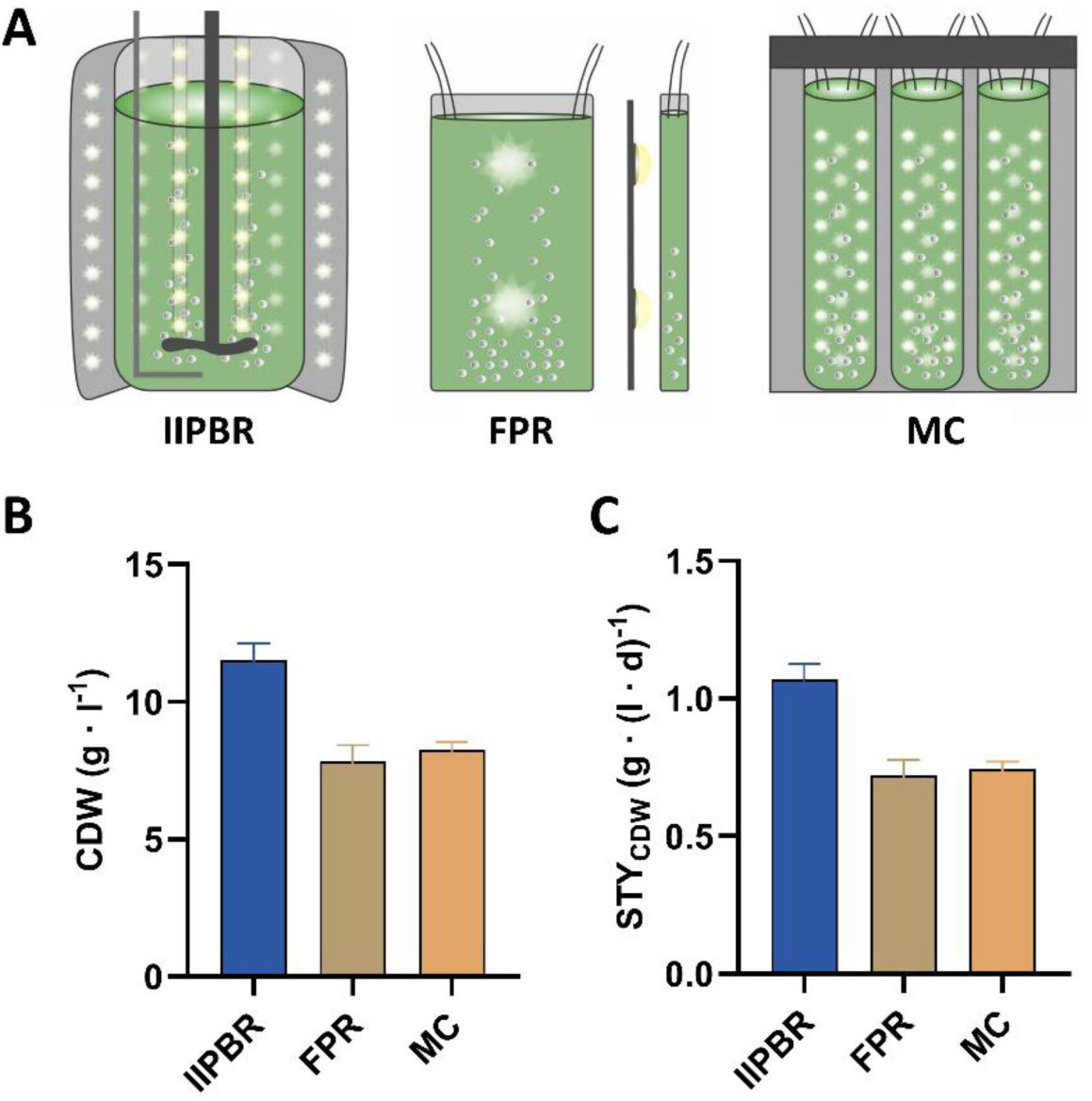
Comparative cultivation of *C. merolae* in different photobioreactor cultivation platforms. (A) Schematic illustration of internally illuminated photobioreactor (IIPBR), flat-panel photobioreactor (FPR) and multi-cultivator (MC). (B) Biomass concentration of *C. merolae* during the cultivation in IIPBR, FPR and MC, calculated form OD values at 750 nm (Figure S1A). Cultivation was carried out for 258 h in the IIPBR, 261 h in the FPR and 266 h in the MC. (C) STY_CDW_ during the cultivations, calculated from the CDW values and cultivation times. The biomass concentration and STY_CDW_ shown for the MC are the values from previous MC cultivation (Figure 1). Cultivations were performed as biological duplicates (IIPBR) or triplicates (FPR and MC).

As shown in Figure 3B and C and summarized in Table 2, the highest biomass yield was reached using the IIPBR. Although exhibiting a slightly lower maximum growth rate of 0.023 ± 0.003 h^-1^ compared to the FPR (0.03 ± 0.002 h^-1^) and MC (0.034 ± 0.001 h^-1^), significantly higher final OD (40.2 ± 2.1) and CDW (11.5 ± 0.6 g l^-1^) values were achieved. Accordingly, the overall STY_CDW_ of 1.07 ± 0.06 g · (l · d)^-1^ in the IIPBR was approximately 1.5-fold higher than in the FPR and MC. Notably, the growth rates obtained in this study exceeded previously reported values of approximately 0.006 h^-1^ and 0.011 h^-1^ described by Rahman et al. (2017) and Krupnik et al. (2023), respectively, but remained below the growth rate of approximately 0.075 h^-1^ reported by Minoda et al. (2004) under further optimized medium conditions. This suggest that additional medium optimization may represent a promising strategy to further enhance the growth performance of *C. merolae* in future studies. When considering the total biomass produced, the advantage of the IIPBR becomes even more pronounced, yielding approximately 5-fold more biomass than the FPR and up to 26-fold more than the MC at the end of the cultivation (Table 2). Although this difference is partly attributable to the larger culture volume of the IIPBR, the system also provides clear advantages in terms of scalability. In particular, the integrated light rods facilitate scale-up by maintaining efficient light distribution throughout the culture volume, even at larger reactor volumes (Figure 3A).

**Table 2:** Growth characteristics of *C. merolae* cultivated in different photobioreactor systems.

| Parameter | IIPBR | FPR | MC |
| --- | --- | --- | --- |
| **t (h)** | 258 | 261 | 266 |
| **$\mu_{\max}$ (h<sup>-1</sup>)** | 0.023 ± 0.003 | 0.03 ± 0.002 | 0.034 ± 0.001 |
| **t<sub>d</sub> (h<sup>-1</sup>)** | 30.9 ± 3.8 | 23 ± 1.2 | 20.4 ± 0.9 |
| **Final OD (-)** | 40.2 ± 2.1 | 27.3 ± 2.1 | 28.8 ± 1 |
| **Final CDW (g · l<sup>-1</sup>)** | 11.5 ± 0.6 | 7.8 ± 0.6 | 8.3 ± 0.3 |
| **STY<sub>CDW</sub> (g · (l · d)<sup>-1</sup>)** | 1.07 ± 0.06 | 0.72 ± 0.06 | 0.75 ± 0.03 |
| **Volume (l)** | 1.6 | 0.45 | 0.08 |
| **Total biomass (g)** | 18.4 ± 0.6 | 3.5 ± 0.05 | 0.7 ± 0.02 |

### Cultivation in internally illuminated photobioreactor combines high biomass and pigment yields

Following the evaluation of growth characteristics, pigment and fatty acid accumulation were analyzed. An overview of the extraction workflow is shown in Figure 4. After cultivation, the biomass was lyophilized and mechanically disrupted. The phycobiliproteins R-PC and R-allophycocyanin (R-APC) were extracted using water as a solvent, while chlorophyll and carotenoids were extracted using methanol. Due to solubility limitations, multiple extraction steps were performed to ensure complete recovery of all compounds. Fatty acids were converted to fatty acid methyl esters (FAMEs) with acidic methanol and subsequently extracted using heptane.

**Figure 4:**
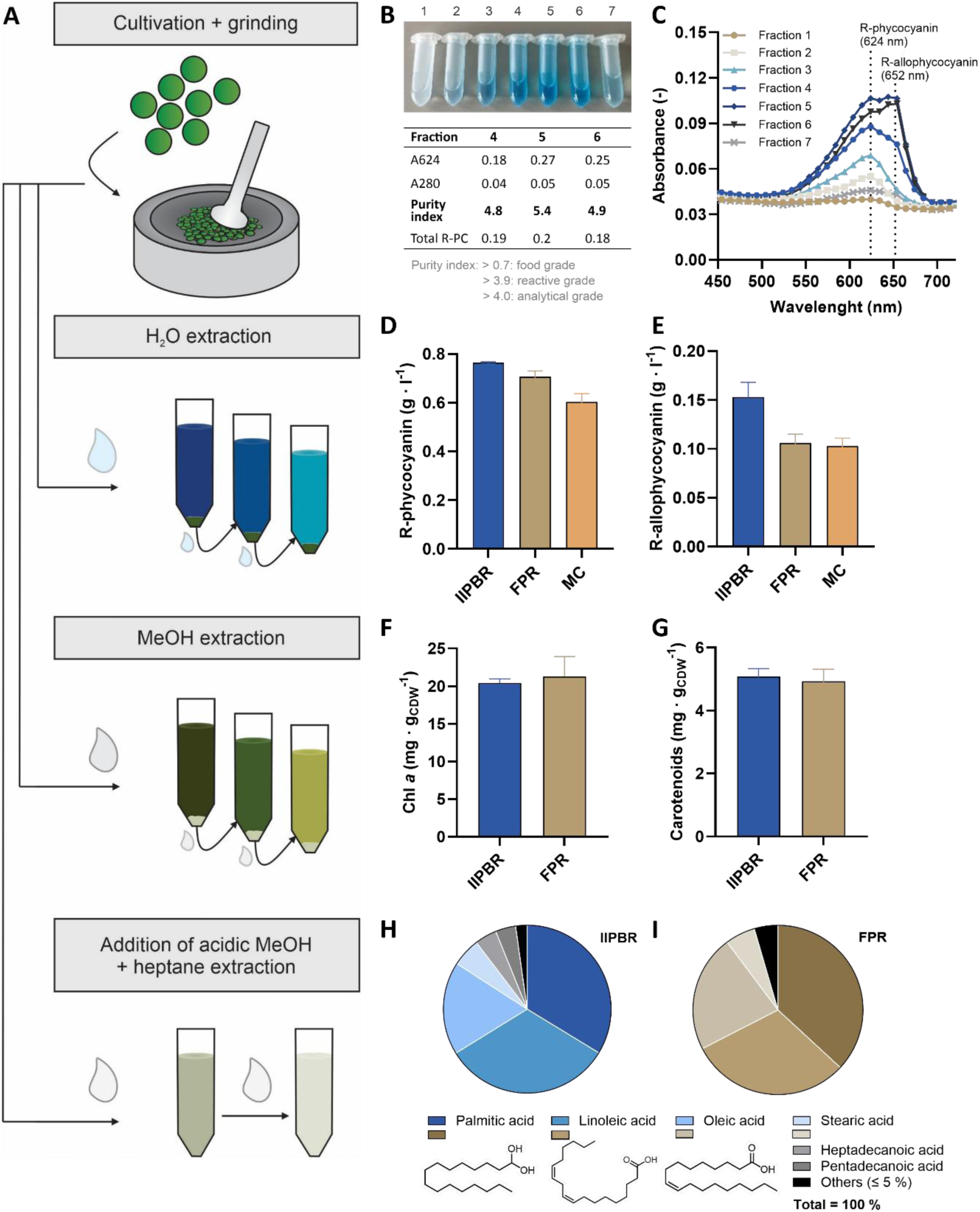
Extraction of pigments and fatty acids from *C. merolae* biomass upon cultivation in different photobioreactors. (A) Extraction scheme for pigment and fatty acid recovery. (B) Elution fractions during purification of R-PC and R-APC from *C. merolae* (2 ml each) via an anion-exchange chromatography and corresponding purity indices, defined as the ratio of absorbance of R-PC to the absorbance of total protein at 280 nm (Patil et al., 2006). (C) Wavelength scans of all ÄKTA fractions for determination of the absorbance maxima for R-PC and R-APC. (D) R-phycocyanin and (E) R-allophycocyanin concentrations as well as (F) Chlorophyll *a* and (G) carotenoids per CDW during cultivation of *C. merolae* in IIPBR, FPR and MC. Concentrations were calculated from an absorption spectrum following bead-based cell disruption and aqueous extraction. Extractions were performed as technical duplicates and cultivations as biological duplicates (IIPBR) or triplicates (FPR and MC). Relative fatty acid profile during cultivation of *C. merolae* in (H) IIPBR and (I) FPR. Structures of major fatty acids are shown below the diagram.

Pigment concentrations can generally be determined using published extinction coefficients. While this approach is reliable for chlorophylls and carotenoids, accurate quantification of R-PC and R-APC requires strain-specific absorbance maxima, as the spectral properties of the chromophore are influenced by its surrounding protein environment (Brejc et al., 1995; Chen et al., 2022; Peng et al., 2014). Many studies rely on absorbance maxima originally established for cyanobacterial systems (Bennett & Bogorad, 1973), although these values may not fully reflect the spectral characteristics of *C. merolae*. This is further supported by reported variability in phycocyanin absorption maxima across different species, including 616 nm for *L. platensis* at pH 5 (Jespersen et al., 2005), 609 nm for the thermophile *Synechococcus lividus* at pH 6 (Edwards et al., 1997), and even values of around 650 nm for *Synechococcus* sp. at pH 3 (Glazer & Fang, 1973), highlighting the strong environmental and species dependence of spectral properties. Rahman et al. (2017) experimentally determined the absorbance maximum of 624 nm for R-PC from *C. merolae*, which was purified using ammonium sulfate precipitation, whereas corresponding organism-specific spectral data for R-APC have not yet been described. To achieve higher-resolution separation and thereby enable determination of the R-APC absorbance maximum, we adapted an anion-exchange chromatography method originally developed for C-PC and C-APC purification from *Synechococcus* sp. PCC6715 to *C. merolae* (Antecka et al., 2025). (Figure S3).

Binding and high-purity elution of the phycobiliproteins were achieved, as confirmed by high purity indices (Figure 4B). With the described procedure, we achieved analytical grade for R-PC. To determine the specific absorbance values, wavelength scans were performed on all elution fractions, and corresponding absorbance maxima were identified directly from the spectral profiles (Figure 4C). This analysis revealed a distinct maximum at 624 nm for R-PC, in agreement with the value reported by Rahman et al. (2017). In contrast to previous work, the higher resolution separation also enabled the detection of a second peak, allowing the identification of an absorbance maximum of 652 nm for R-APC. In addition, the elution profiles showed that R-PC eluted at lower salt concentrations (Fractions 1-3), whereas R-APC eluted at higher salt concentrations (Fractions 4-7), indicating that this method also enables partial separation of the phycobiliproteins.

With the experimentally determined absorbance maxima, R-PC and R-APC concentrations were quantified in the biomass obtained from the different cultivation platforms (Figure 4D and E). The R-PC concentration was highest in the IIPBR compared to the FPR and MC. In contrast, the R-PC content per biomass was similar across all cultivation platforms, ranging between 0.07 and 0.09 g_R-PC_ · g_CDW_^-1^, indicating that under the chosen conditions R-PC production is primarily dependent on biomass concentration rather than the cultivation system itself (Table 3). Similar trends were observed for R-APC, although its overall concentration was consistently lower than that of R-PC (Figure 4E and Table 3).

**Table 3:** Pigment concentrations of *C. merolae* cultivated in different photobioreactor systems.

| Parameter | IIPBR | FPR | MC |
| --- | --- | --- | --- |
| **Final R-PC concentration (g · g<sub>CDW</sub><sup>-1</sup>)**<br>**/ Total R-PC (g)** | 0.07 ± 0.003<br>**/ 1.29 ± 0.06** | 0.09 ± 0.01<br>**/ 0.32 ± 0.04** | 0.07 ± 0.005<br>**/ 0.05 ± 0.00** |
| **Final R-APC concentration (g · g<sub>CDW</sub><sup>-1</sup>)**<br>**/ Total R-APC (g)** | 0.013 ± 0.005<br>**/ 0.24 ± 0.09** | 0.014 ± 0.002<br>**/ 0.05 ± 0.00** | 0.012 ± 0.001<br>**/ 0.01 ± 0.00** |
| **Final Chl *a* concentration (mg · g<sub>CDW</sub><sup>-1</sup>)**<br>**/ Total Chl *a* (mg)** | 20.4 ± 0.4<br>**/ 375.4 ± 7.4** | 21.3 ± 2.2<br>**/ 74.6 ± 7.7** |  |
| **Final carotenoids concentration (mg · g<sub>CDW</sub><sup>-1</sup>)**<br>**/ Total carotenoids (mg)** | 5.1 ± 0.2<br>**/ 93.8 ± 3.7** | 4.9 ± 0.3<br>**/ 17.2 ± 1.1** |  |
| **Total biomass (g)** | **18.4 ± 0.6** | **3.5 ± 0.05** | **0.7 ± 0.02** |

In addition to R-PC and R-APC, other photopigments such as chlorophyll *a* (Chl *a*) and carotenoids are of industrial interest, particularly for applications in the food and feed sectors (Oliveira et al., 2024). Accordingly, Chl *a* and carotenoids were extracted from biomass of the different cultivation systems and showed comparable concentrations in the IIPBR and FPR, with approximately 20 mg g_CDW_^-1^ Chl a and 5 mg g_CDW_^-1^ carotenoids (Figure 4F and G, Table 3).

Although pigment concentrations were similar, the IIPBR yielded the largest total pigment amounts due to its higher final biomass concentration, supported by the superior scalability of the IIPBR (Table 3). Specifically, the IIPBR produced approximately 4-fold more R-PC than the FPR and 26-fold more than the MC. For R-APC, increases of approximately 5-fold and 24-fold were observed compared to the FPR and MC, respectively (Table 3). Similarly, total Chl a and carotenoids yields were approximately 5-fold higher in the IIPBR than in the FPR.

Beyond pigments, the fatty acid profile of *C. merolae* is of interest for applications such as cosmetics. The predominant fatty acids detected were palmitic acid (C16:0, 35 %), linoleic acid (C18:2 Δ9,12c, 31 %) and oleic acid (C18:1 Δ9c, 20 %), consistent with previous reports (Fukuda et al., 2018). Stearic acid (C18:0) was also present with approximately 5 %. The relative abundance of these major fatty acids was similar across all cultivation systems (Figure 4H and I and S4), indicating that reactor configuration had only a minor influence on the overall fatty acid profile. However, distinct differences were observed in minor components, as heptadecanoic acid (C17:0) and pentadecanoic acid (C15:0), each accounting for approximately 4 % of total fatty acids, were detected exclusively in the IIPBR. Additional minor fatty acids (below 5 %) included myristic acid (C14:0), γ-homolinoleic acid (C20:3 Δ8,11,14c), gondoic acid (C20:1 Δ11c) and cis-10-heptadecenoic acid (C17:1 Δ10c).

### High protein and starch content of *C. merolae* supports its potential use as renewable feedstock for heterotrophs

After identifying the IIPBR as the most effective cultivation platform for both biomass and pigment production, the resulting *C. merolae* biomass was further characterized by elemental analysis. Since cultivation was performed under phototrophic conditions, the carbon (C) content provides an indication of CO_2_ fixation efficiency. The elemental analysis revealed comparable C-contents for biomass obtained from the IIPBR (47.1 ± 0.2 %) and FPR (46.8 ± 0.5 %) (Figure 5A), whereas the MC exhibited a slightly lower C-content (45.2 ± 0.6 %) (Figure S4). These results are in line with other reports showing a C-content up to 50 % for *C. merolae* (Dandamudi et al., 2020; Muppaneni et al., 2017). Under identical conditions, *G. javensis* and *L. platensis* reached C-contents of 40.9 ± 0.1 % and 30.0 ± 0.4 %, respectively (Figure S5), indicating significantly lower CO_2_ fixation efficiencies compared to *C. merolae*. In contrast, the hydrogen (H) contents were comparable across all species and cultivation platforms, whereas nitrogen (N) contents were 3-fold and 2-fold higher in *C. merolae* as for *G. javensis* and *L. platensis*, respectively (Figure S5). Consequently, the calculated carbon-to-nitrogen (C:N) ratios of *C. merolae* (5.4 (IIPBR), 6.8 (FPR), and 4.5 (MC)) were consistently lower than those of *G. javensis* (approximately 13 across the cultivation platforms). This suggests a larger proportion of carbohydrates in *G. javensis*, likely associated with its thick and rigid cell wall. *L. platensis* showed comparable C:N ratios across the cultivation platforms, indicating a similar balance between protein content and carbon storage compounds. Oxygen was not directly measured in the CHN analysis, however, the unknown fraction (37.2 %) is likely attributed to oxygen, as reported elemental compositions of *C. merolae* indicate an oxygen content of approximately 36.5 % (Li et al., 2021).

**Figure 5:**
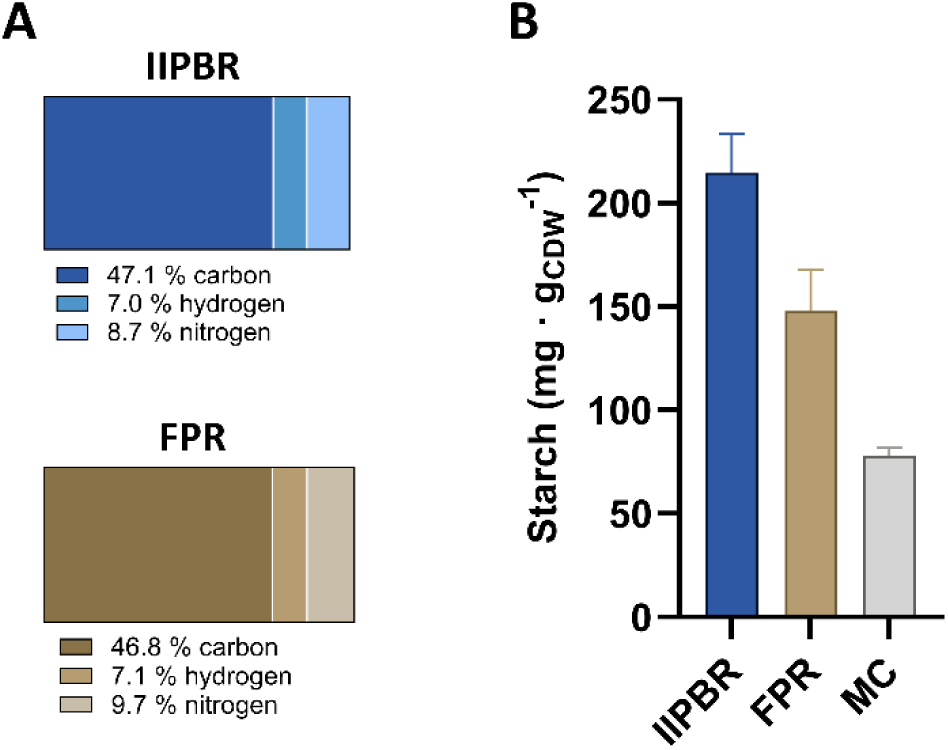
Elemental analysis and starch content of the endpoint samples. (A) Elemental analysis and (B) starch content. Extractions were performed as technical triplicates and cultivations as biological duplicates (IIPBR) or triplicates (FPR).

The starch content of *C. merolae* cultivated in the IIPBR accounted for 21.5 ± 1.7 % of the cell dry weight (215 ± 17 mg g_CDW_^-1^), exceeding the values observed in the FPR (14.8 ± 1.6 %; 148 ± 16 mg g_CDW_^-1^) and the MC (7.8 ± 0.3 %; 78 ± 3 mg g_CDW_^-1^) (Figure 5B). This indicates that the IIPBR provides favorable conditions for intracellular carbon storage, consistent with the elemental analysis showing the highest carbon fixation efficiency in this cultivation platform. The observed starch levels are in agreement with previously reported values for microalgae under nutrient-replete conditions, where storage α-glucans such as floridean starch represent a major carbohydrate fraction. In contrast, the lower starch accumulation in the FPR and MC suggests suboptimal conditions for carbon assimilation or storage under these cultivation settings. Importantly, the combination of high protein content and substantial starch accumulation highlights the potential of *C. merolae* biomass as feedstock for heterotrophs, as starch represents a readily accessible carbon source while proteins supply nitrogen-rich nutrients required for microbial growth (Miao et al., 2018; Wang et al., 2020). Although the starch content observed in the IIPBR already demonstrates considerable biotechnological potential, previous studies showed that starch accumulation in *C. merolae* can be further enhanced under nitrogen depletion or inhibition of the target of rapamycin (TOR) pathway using rapamycin, with TOR inactivation reported to induce up to a 10-fold increase in starch accumulation (Pancha et al., 2019). In addition, metabolic engineering approaches targeting TOR-regulated starch synthesis pathways, including modification of glycogenin-like proteins such as CmGLG1, were also shown to enhance starch accumulation. However, such conditions are typically accompanied by reduced protein content and impaired growth performance. Further studies could therefore explore cultivation strategies and metabolic engineering approaches to optimize the balance between starch accumulation, protein content, and overall biomass productivity, thereby tailoring biomass composition to the respective application.

### Monosaccharide and glycosidic linkage analysis indicate structurally complex cell wall composition

Although various metabolites and biotechnological applications of *C. merolae* have been investigated, detailed studies of its cell wall polysaccharide matrix remain scare. Hence, monosaccharide and glycosidic linkage analysis was conducted on cell wall preparations to gain more insights into potential enzymatic breakdown strategies of algae biomass (Figure 6).

**Figure 6:**
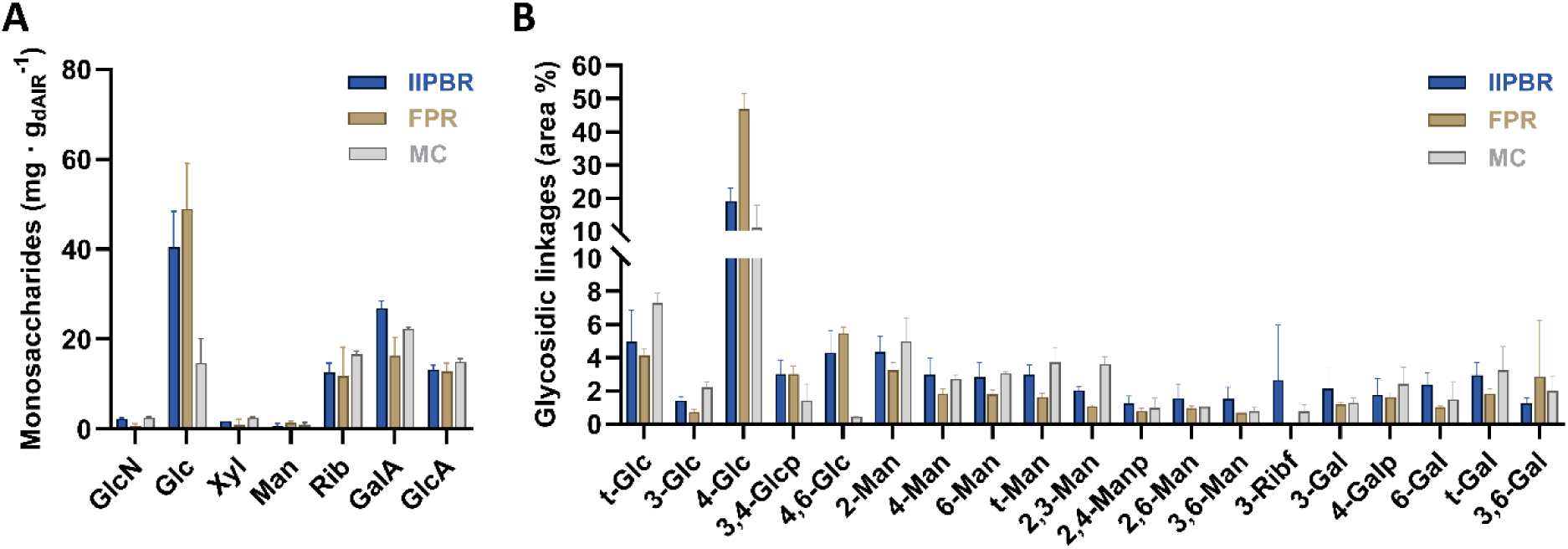
Glycosidic linkage analysis of neutral monosaccharides present in destarched cell wall materials from *C. merolae* under three different cultivation conditions. (A) GlcN – glucosamine; Glc – glucose; Xyl – xylose; Man – mannose; Rib – ribose; GalA – galacturonic acid and GlcA – glucuronic acid. (B) Glc – glucose; Glcp – glucopyranose; Man – mannose; Manp – mannopyranose; Ribf – ribofuranose; Gal – galactose; Galp – galactopyranose. Non-carbohydrate peaks accounted for 34.5 ± 5.4 %, 19.2 ± 2.1 % and 45.4 ± 8.7 % of the total peak area in the IIPBR, FPR and MC, respectively. Cultivations were performed as biological duplicates (IIPBR) or triplicates (FPR).

The detected fraction of cell wall polysaccharides in the destarched, alcohol-insoluble residue (dAIR) was comparatively low (approximately 6 %) but structurally complex (Figure 6A). The total monosaccharide content varied across the different cultivation platforms, with the highest values detected in the IIPBR (96.6 ± 2.2 mg g_dAIR_^-1^) and the FPR (92.9 ± 2.3 mg g_dAIR_^-1^), followed by the MC (74.8 ± 3.3 mg g_dAIR_^-1^). The monosaccharide composition also differed between the systems. In both the IIPBR and FPR, glucose was the dominant monosaccharide (approximately 50 %), followed by galacturonic acid (GalA), glucuronic acid (GlcA), whose abundance indicates polyanionic polymers such as pectic polysaccharides in higher plants, and ribose. In contrast, biomass from the MC was dominated by GalA, while glucose was present only in minor amounts (14.7 ± 3.8 mg g_dAIR_^-1^). These differences suggest that the cultivation platforms influence not only total carbohydrate content, but also the relative distribution of structural polysaccharides such as glucans vs polyanionic polysaccharides within the cell wall. Overall, the carbohydrate fraction of *C. merolae* was primarily composed of glucose, ribose, GalA and GlcA, differing from red microalgae such as *Porphyridium* spp. and *Dixoniella griesa*, which have been reported to contain cell wall polysaccharides enriched in glucose, xylose and galactose (Arad & Levy-Ontman, 2010). Minor amounts of glucosamine, xylose and mannose were detected across all systems, indicating the presence of additional, less abundant polysaccharide components, which is consistent with observations in the related species *Cyanidium caldarium* (Bailey & Staehelin, 1968). In addition, *C. merolae* cultivated in shake flasks under 0.04 % CO_2_ showed an expanded monosaccharide profile, including Rhamnose, Arabinose and galactose, further supporting that the cell wall polysaccharide composition is strongly dependent on the cultivation platform and conditions. This variability is particularly relevant for the use of *C. merolae* biomass as third generation feedstock, as consistent substrate composition is important for efficient and predictable microbial metabolization.

An extended glycosidic linkage analysis of the neutral monosaccharides present in the cell wall material revealed that 1,4-linked glucopyranosyl residues (4-Glc) constitute the by far predominant structural motif indicating the presence of cellulose (Figure 6B). Cellulose was present in all cultivation systems, with the highest proportion observed in the FPR, followed by the IIPBR, and the MC.

However, in addition to cellulose-derived glucopyranosyl linkages, several other glucopyranosyl residues were detected, including terminal Glc, 3-substituted Glc, 3,4-Glcp, 4,6-Glc, suggesting the abundance of branched glucan structures. Furthermore, the detection of structurally diverse mannopyranosyl and galactopyranosyl linkages, including branched configurations such as 2,3-, 2,6-, 3,6-linked Man as well as 3,6-linked Gal, suggests a complex cell wall architecture. In particular the mannosyl-linkages are reminiscent of mannoproteins. The variety of glycosidic linkage patterns indicates the presence of structurally diverse glucan-, galactan– and mannan-containing polysaccharides or glycoconjugates within the cell wall matrix. Overall, these findings suggest that the cell wall of *C. merolae* is more structurally complex than previously assumed and potentially comprise polysaccharide classes comparable to those described in other unicellular red algae, including cellulose and polyanionic pectin-like structures, branched glucans, sulfated polysaccharides and glycoproteins (Arad & Levy-Ontman, 2010; Bailey & Staehelin, 1968; Geresh et al., 1990). Overall, the presented results demonstrate that *C. merolae* exhibits not a relatively simple cellular architecture with only minimal cell wall structures but harbors a distinctive biomass composition, characterized by high protein and starch contents, alongside structurally diverse cell wall carbohydrates that are responsive to cultivation conditions.

## Conclusion

In this study, *C. merolae* was systematically evaluated across different photobioreactor systems, addressing the current lack of comparative data on biomass productivity and biochemical composition under varying reactor conditions. Among the tested platforms, the IIPBR proved to be the most effective, consistently reaching the highest biomass yield. The resulting biomass was characterized by high nitrogen (8.7 %) and carbon (47.1 %) content, as well as by the accumulation of large amounts of valuable compounds, including phycobiliproteins, pigments and fatty acids. In addition, the substantial accumulation of starch within the cytoplasm (21.5 %) highlights its potential as a carbon source for heterotrophic processes. Despite a comparatively low fraction of cell wall polysaccharides, their structurally diverse composition and sensitivity to cultivation conditions underline an additional level of cell wall synthesis adaptability.

Overall, the results demonstrate that the cultivation platform has a decisive impact on both biomass formation and metabolic profiles. Together, these findings advance our understanding of reactor-dependent physiology in extremophilic microalgae and provide an important basis for optimizing scalable bioprocesses, thereby supporting the establishment of *C. merolae* as a versatile chassis for biotechnological applications.

## Material and Methods

### Chemicals and strains

The chemicals used in this study were obtained from Sigma-Aldrich (St. Louis, USA), Thermo Fisher Scientific (Waltham, USA) or VWR (Radnor, USA) and were of analytical grade. The strains used in this study were *C. merolae* 10D (NIES-3377) (De Luca et al., 1978), *G. javensis* 074G (NIES-3638) (Ott & Seckbach, 1994) and *L. platensis* PCC7345 (DSM-25777) (Gomont, 1893), which were kindly provided by Prof. Dr. Andreas Weber from Heinrich Heine University Düsseldorf, Germany.

### Media and culture conditions

Maintenance cultures were inoculated monthly for long-term storage. *C. merolae*, *G. javensis*, and *L. platensis* were cultivated in MA2 medium (Allen, 1959), 2xGS modified Allen medium (Allen, 1959) and Spirulina medium (Aiba & Ogawa, 1977), respectively, at an initial OD_750_ or OD_680_ of 0.1 in 50 mL medium in 100 mL baffled shake flasks. Cultures were maintained at room temperature on a laboratory bench without additional gassing or illumination, and agitated at 20 rpm using an orbital shaker (IKA Rocker 3D Digital, Germany).

Precultures for main cultivations were inoculated from maintenance cultures into 100 mL medium in 500 mL baffled shake flasks and incubated at 42 °C (*C. merolae*, *G. javensis*) or 30 °C (*L. platensis*) at 100 rpm in a shaker incubator (Infors HT, Switzerland). Cultures were illuminated with 100 µE using white LED panels (Paros, Netherlands) without additional gassing.

Small-scale cultivations were performed in vertical column photobioreactors using a Multi-Cultivator MC-1000 OD (Photon Systems Instruments, Czech Republic). The device was equipped with an additional gas disperser pipe connected to a CO_2_ Controller 2000 (Pecon, Germany), enabling cultivation with CO_2_-enriched air. *C. merolae*, *G. javensis* and *L. platensis* were inoculated from precultures to an OD of 0.5 and cultivated at 42 °C (*C. merolae* and *G. javensis*) or 30 °C (*L. platensis*) under continuous illumination of 50-700 µE. Cultures were aerated with 5 % (v v^-1^) CO_2_-enriched air at a flow rate of 10.6 sl h^-1^. Microfluidic cultivation experiments were conducted using an inverted microscope (Ti-E, Nikon, Japan), that was upgraded for phototrophic time-lapse cultivation experiments. For a detailed description of the platform used for microfluidic cultivation experiments, the production of the silicon wafer by direct laser writing and PDMS soft lithography, the reader is referred to Witting et al. (2025). In short: A heating incubator (Incubator: NL 2000, Temperature controller: Temp-Controller 2000-2, PECON, Germany) ensured a stable temperature (42 °C) throughout the cultivations. Cold white illumination (30 µE m^-2^ s^-1^) to drive photosynthesis was generated by an external light engine (Spectra Tune Lab, LEDMOTIVE, Spain) coupled into a ring light (A08360, Schott, Germany) fixed around the microscope condenser. Growth medium was continuously supplied at 200 nl min^-1^ via a syringe pump (Nemesys, CETONI, Germany). For soft lithography PDMS and curing agent (SYLGARD 184) were mixed in a 10:1 ratio, degassed, poured over the master-mold, heated for 2 h at 80 °C, and cut into appropriate pieces. Chips were washed for 45 min in n-pentane, and subsequently for 45 min in acetone twice. Finally, inlet and outlet holes were created and the cultivation chip was bonded onto a glass substrate (D263®Bio, Schott, Germany) by plasma bonding (Duration: 25 s, Evacuation pressure: 0.16 mbar, Plasma power 510 W). Samples were injected into the cultivation chip and positions for imaging (1 per hour) were selected. Cell counts per frame were determined through manual counting.

Large-scale cultivations were performed in a custom-designed, internal-illuminated photobioreactor based on the DASGIP system (Eppendorf, Germany). The reactor (total volume: 2.3 L; working volume: 1.6 L) was equipped with two Rushton turbines, a propeller stirrer, an external illumination module, and internally illuminated glass rods. *C. merolae* was cultivated in MA2 medium and inoculated at OD 0.5. Cultivation was performed at 42 °C, with agitation at 200 rpm (0-168 h), followed by 400 rpm until the end. Aeration was performed with 5 % CO_2_ and 95 % N_2_ at 20 sl h^-1^ via an aeration stone. The light intensity was gradually increased over the cultivation time from 70 µE to 1100 µE. Flat-panel photobioreactors (25 x 25 x 1 cm, total volume: 550 mL) were used as an additional larger-scale cultivation platform (Forschungszentrum Jülich, IMET). *C. merolae* was inoculated at OD 0.5 into 450 mL MA2 medium and cultivated at approximately 42 °C. Illumination was provided from both sides using 60 W, 700 lumen bulbs (Philips, Netherlands; Leuci, Germany). Cultures were aerated with 5 % CO_2_ and 95 % N_2_ at 20 sl h^-1^ using a DASGIP MF4 module (Eppendorf, Germany) or red-y gas flow controllers with the corresponding control software (Vögtlin, Switzerland).

Growth in all cultivations, except for microfluidic growth chambers, was monitored by offline OD measurements. Samples (3-5 mL) were collected and centrifuged in 1 mL aliquots at 1000 x g for 45 min (*C. merolae*) or 6000 x g for 10 min (*G. javensis* and *L. platensis*) at 4 °C. The resulting cell pellets were stored at –20 °C until further analysis. After reaching the final OD, cells were harvested by centrifugation, washed with pre-cooled 2.7 % (w/v) NaCl solution, and subsequently freeze-dried (LyoCube, Christ, Germany).

### Purification of R-PC and R-APC

R-PC and R-APC were purified following a workflow adapted from a previously published protocol (Antecka et al., 2025). Lyophilized *C. merolae* cells were lysed by two freeze-thaw cycles in 10 mM sodium acetate buffer (pH 5.0), followed by centrifugation for removal of cell debris. The resulting supernatant was subjected to anion-exchange chromatography. Purification was performed using a 1 mL prepacked Q Sepharose Fast Flow (Q FF) column on an ÄKTA Pure fast protein liquid chromatography (FPLC) system (GE Healthcare). The column was equilibrated with 20 column volumes (CV) of 10 mM sodium acetate buffer (pH 5.0). After sample application, unbound material was removed by washing with equilibration buffer. Bound proteins were eluted using a two-step linear gradient of NaCl in 10 mM sodium acetate buffer (pH 5.0): 0-180 mM over 10 CV, followed by 180-1000 mM over an additional 10 CV, at a flow rate of 1 mL min^-1^. Elution was monitored by measuring absorbance at 280 nm and conductivity, and fractions were collected in 2 mL volumes. Following separation, fractions were stored at 4 °C and subsequently analysed by wavelength scanning to identify R-PC– and R-APC-containing fractions as well as the specific absorbance maxima.

### Extraction and quantification of pigments and fatty acids

To determine R-PC and R-APC concentrations in *C. merolae*, frozen biomass samples were resuspended in 1 mL pre-cooled phosphate-buffered saline (PBS). Cell disruption was performed using a Precellys24 homogenizer (Bertin Technologies, France), with 200 µL of 0.1 mm glass beads in safe-lock tubes. Samples were homogenized three times at 4000 rpm for 30 s. Following homogenization, samples were incubated on ice with gentle shaking for 60 min, and subsequently centrifuged at 16000 x g and 4 °C for 60 min. The clarified supernatant was used for absorbance measurements. Absorption spectra were recorded using an Infinite M1000 Pro microtiter plate reader (Tecan, Switzerland), and R-PC and R-APC concentrations were calculated using published spectrophotometric equations with previously determined specific absorbance maxima.

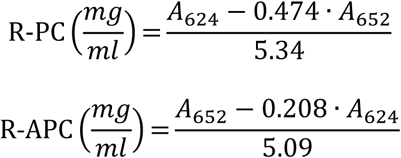

For chlorophyll *a* and carotenoid analysis, frozen samples were resuspended in 1 mL pre-cooled methanol. Cell disruption was performed as described above, using the Precellys24 system. After homogenization, samples were incubated on ice for 30-60 min with gentle shaking. Cell debris was removed by centrifugation at 16000 x g and 4 °C for 60 min. The clarified supernatant was used for absorbance measurements as described above. Chl a and carotenoid concentrations were calculated using established equations.

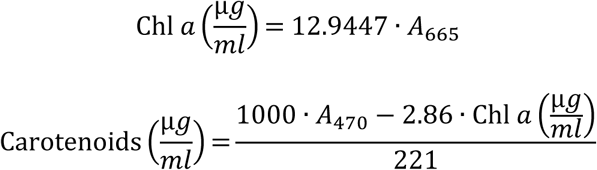

The fatty acid composition was determined by gas chromatography coupled to time-of-flight mass spectrometry (GC-ToF-MS). Approximately 10 mg of lyophilized biomass was transferred into Pyrex glass tubes and resuspended in 1.7 mL acidic methanol containing 10 % (v v^-1^) sulfuric acid for transesterification. Samples were incubated at 60 °C for 4 h at 750 rpm in a Thermomixer (Eppendorf, Germany). After cooling to room temperature, 300 µL ddH_2_O and 1.2 mL heptane were added. Following vigorous mixing, the heptane phase containing fatty acid methyl esters (FAMEs) was separated and stored at –20 °C until analysis. FAMEs were analysed using a method previously described (Morschett et al., 2019). Fatty acid composition was calculated as the relative proportion of all identified fatty acids.

### Elemental analysis, starch quantification and cell wall polysaccharide analysis

The elemental composition was determined using a Vario EL CUBE analyzer (Elementar, Langenselbold, Germany). Approximately 5 mg of lyophilized biomass was analysed in the CHN mode to quantify the carbon (C), hydrogen (H) and nitrogen (N) content.

The starch content of *C. merolae* was determined gravimetrically via quantification of the AIR and dAIR, following previously described protocols with minor modifications. Briefly, 10-20 mg of lyophilized biomass was homogenized in 2 mL screw-cap tubes containing two steel beads, using a mixer mill (MM400, Retsch, Germany) for 10 min at 30 Hz. Subsequent AIR preparation, starch removal, and dAIR recovery were performed as described previously by Foster et al. (2010). The monosaccharide composition of dAIR was analysed using high-performance anion-exchange chromatography with pulsed amperometric detection (HPAEC-PAD), as described previously by Robert et al. (2021). For quantification, 100 µL of a 1 mg ml^-1^ fucose solution was added as an internal standard to the dAIR samples and calibration standards. Sugar standards included rhamnose, arabinose, glucosamine, glucose, galactose, xylose, mannose, galacturonic acid and glucuronic acid at concentrations ranging from 0 to 300 µg ml^-1^. Monosaccharides were identified based on retention times of standards, and peak areas were integrated for quantitative analysis. Glycosidic linkage analysis of the dAIR fraction was performed by derivatizing the sample to their partially methylated alditol acetates followed by gas-chromatography electron impact quadrupole mass spectrometry (GC-MS) as described by Liu et al. (2015). The GC-MS system consisted of an Agilent 7890B GC System equipped with a 5977A MSD and an SP-2380 Fused Silica Capillary Column (30 m × 0.25 mm i.d. × 20 μm film thickness, Supelco, USA). The components were identified and quantified based on the retention times of glycosidic linkage standards and their corresponding mass spectra utilizing a spectral database for partially methylated alditol acetates (https://www.ccrc.uga.edu/specdb/ms/pmaa/pframe.html).

## Abbreviations

R-PC: R-phycocyanin; R-APC: R-allophycocyanin; Chl *a*: Chlorophyll a; AIR: alcohol-insoluble residue; MC: multi-cultivator photobioreactor; IIPBR: Custom-designed internal-illuminated photobioreactor; FPR: flat-panel photobioreactor.

## Declarations / Acknowledgements

### Author contributions

JF, MF and PE designed and supervised the study. PE established the phototrophic cultivation platforms and performed the ÄKTA purification. JV performed the cultivations and extracted the pigments. MD performed additional MC cultivations. WL performed the carbohydrate analysis. LW performed and analyzed the microfluidic cultivations. JG performed the GC-ToF-MS analysis. PE, JV and MP analyzed the data. PE drafted the manuscript. JF, MF, PE, VU, DK and MP revised the manuscript and prepared the final version of the manuscript. All authors read and approved the final manuscript.

### Funding

This project has received funding from the Ministry of Culture and Science of the German state of North Rhine-Westphalia (Project: “Verbundprojekt: Active Carbon Capture for Sustainable Synthesis”, Förderkennzeichen: PB22_028B) and the German Research Foundation (CRC 1535, project ID 458090666).

### Availability of data and materials

All data generated or analyzed during this study are included in this published article and its supplementary information files.

## Acknowledgements

We thank Jens Kruse for performing the elemental analysis. We thank Alexander Lehnert for supporting the ÄKTA purification. We thank Daniel Klein, Florian Walkenbach and Jürgen Paulzen for constructing the illumination module. We also thank Silas Beckmann and Melina Gentges for supporting the pigment extractions. In addition, we thank Katharina Grosche for excellent technical assistance performing the monosaccharide and glycosidic linkage analysis. Finally, we thank all project partners for fruitful discussions.

## Conflict of interest

The authors declare that they have no competing interests.

## Notes

### Competing Interest Statement

The authors have declared no competing interest.

