## Supplementary Information, Figure S1-5 for "Systematic evaluation of *Cyanidioschyzon merolae* across photobioreactor systems: Linking reactor design to biomass production and biochemical composition"

**Content:**

Figure S1: Calibration curves for relationship between optical density and cell dry weight of the three microalgae strains tested (n=2).

Figure S2: Growth rates at different light intensities and $CO_2$ concentrations in the MC.

Figure S3: Purification of R-PC and R-APC from *C. merolae*.

Figure S4: Fatty acids and elemental analysis of *C. merolae* endpoint samples.

Figure S5: Elemental analysis of *G. javensis* and *L. platensis* endpoint samples.

Video S1: Time-lapse image sequence displaying growth of *C. merolae* in a microfluidic cultivation chamber.

Video S2: Time-lapse image sequence displaying growth of *G. javensis* in a microfluidic cultivation chamber.

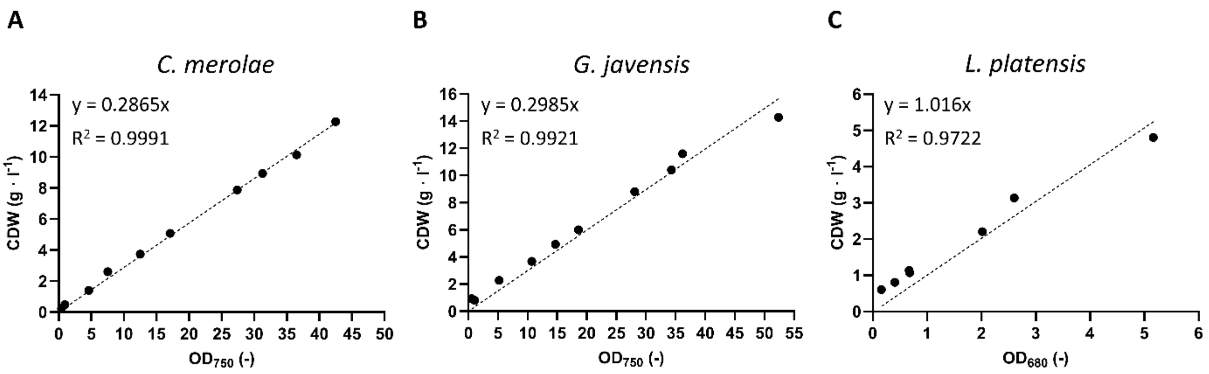


**Figure S1: Calibration curves for relationship between optical density and cell dry weight of the three microalgae strains tested (n=2).**

The linear equations were used to determine the cell dry weights from the optical density values of the cultivation samples (Figure 1 and Table 1).

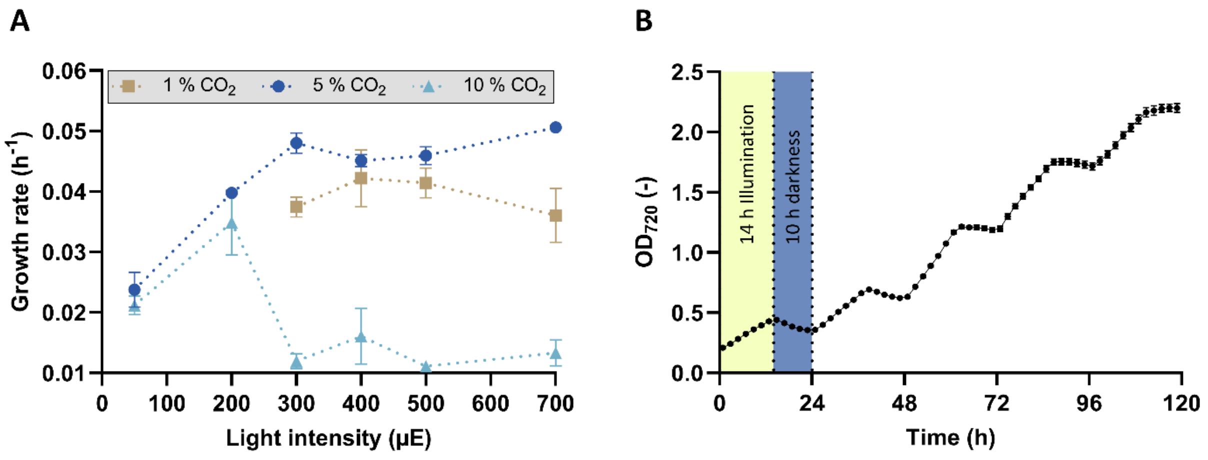


**Figure S2: Growth rates at different light intensities and $CO_2$ concentrations in the MC.**
*C. merolae* 10D day/night cycles (14/10 h). The mean values with standard deviation of two independent biological replicates are shown (n=2).

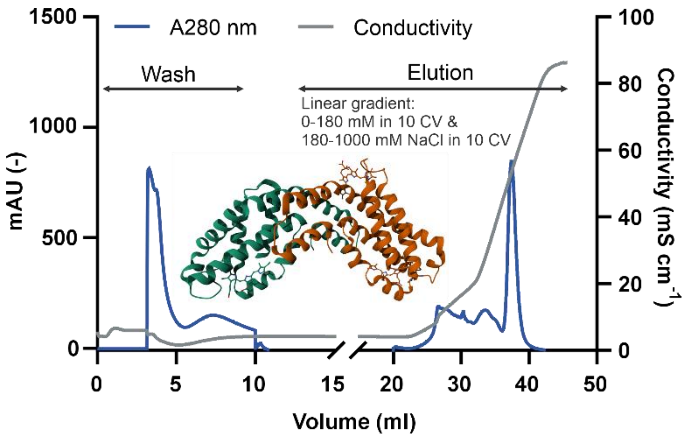


**Figure S3: Purification of R-PC and R-APC from *C. merolae*.**
Chromatogram showing protein elution monitored at 280 nm alongside conductivity. After removal of unbound material, target proteins were eluted using a linear gradient of sodium chloride. PDB code: 1I7Y.

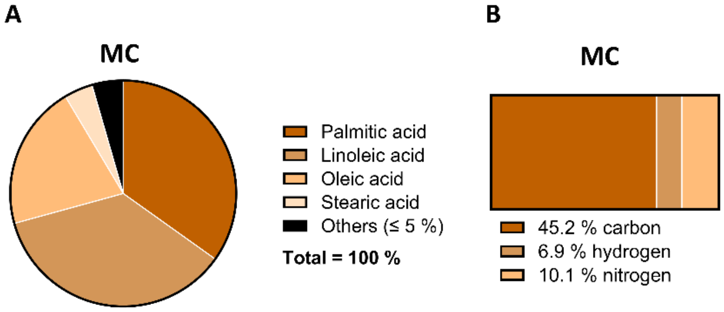


**Figure S4: Fatty acids and elemental analysis of *C. merolae* endpoint samples.**
Fatty acids and (B) elemental analysis of the endpoint samples after cultivation of *C. merolae* in the MC (n=3).

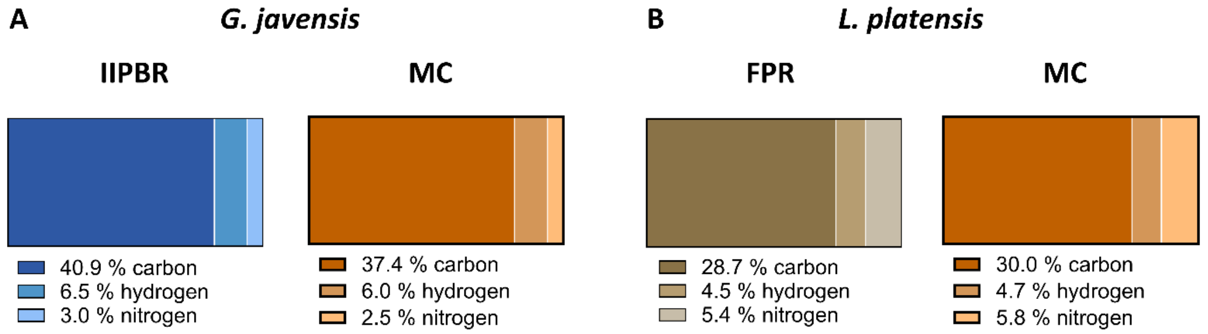


**Figure S5: Elemental analysis of *G. javensis* and *L. platensis* endpoint samples.**
Elemental analysis of (A) *G. javensis* endpoint samples of IIPBR (n=2) and MC (n=3) and (B) *L. platensis* endpoints samples of FPR (n=1) and MC (n=3).

**Video S1: Time-lapse image sequence displaying growth of *C. merolae* in a microfluidic cultivation chamber (same as Figure 2A).** Each image shows an overlay of phase-contrast and fluorescence images (excitation: 514/30 nm; emission: 561 nm; emission: 629/56 nm) acquired during time-lapse microscopy. Images were taken every hour. During the cultivation, the chip was continuously illuminated with 30 µE white light. The temperature was maintained at 42 °C and the growth medium was perfused continuously at a rate of 200 $nl \cdot min^{-1}$. The scale bar indicates a length of 2 µm.

**Video S2: Time-lapse image sequence displaying growth of *G. javensis* in a microfluidic cultivation chamber (same as Figure 2B).** Each image shows an overlay of phase-contrast and fluorescence images (excitation: 514/30 nm; emission: 561 nm; emission: 629/56 nm) acquired during time-lapse microscopy. Images were taken every hour. During the cultivation, the chip was continuously illuminated with 30 µE white light. The temperature was maintained at 42 °C and the growth medium was perfused continuously at a rate of 200 $nl \cdot min^{-1}$. The scale bar indicates a length of 2 µm.
